# A de novo genome assembly and annotation of the southern flying squirrel (*Glaucomys volans*)

**DOI:** 10.1101/2021.06.09.447775

**Authors:** J.F. Wolf, J. Bowman, S. Keobouasone, R.S. Taylor, P.J. Wilson

**Affiliations:** Biology Department, Trent University, 1600 West Bank Drive, Peterborough, ON K9J 7B8, Canada; Ontario Ministry of Natural Resources and Forestry, Wildlife Research and Monitoring Section, Trent University, DNA Building, 1600 West Bank Drive, Peterborough, ON K9J 7B8, Canada; Landscape Science and Technology Division, Environment and Climate Change Canada, 1125 Colonel By Drive, Ottawa, ON K1S 5R1, Canada

**Keywords:** Northern flying squirrel, Southern flying squirrel, *Glaucomys volans*, *Glaucomys sabrinus*, hybrid zone, introgression, comparative genomics

## Abstract

Northern (*Glaucomys sabrinus*) and southern (*Glaucomys volans*) flying squirrels are widespread species distributed across much of North America. Northern flying squirrels are common inhabitants of the boreal forest, also occurring in coniferous forest remnants farther south, whereas the southern flying squirrel range is centered in eastern temperate woodlands. These two flying squirrel species exhibit a hybrid zone across a latitudinal gradient in an area of recent secondary contact. *Glaucomys* hybrid offspring are viable and can successfully backcross with either parental species, however, the fitness implications of such events are currently unknown. Some populations of *G. sabrinus* are endangered, and thus, interspecific hybridization is a key conservation concern in flying squirrels. We sequenced and assembled a *de novo* long-read genome from a *G. volans* individual sampled in southern Ontario, Canada, while four short-read genomes (2 *G. sabrinus* and 2 *G. volans*, all from Ontario) were re-sequenced on Illumina platforms. The final genome assembly consisted of approximately 2.40Gb with a scaffold N50 of 455.26Kb. Benchmarking Universal Single-Copy Orthologs reconstructed 3,742 (91.2%) complete mammalian genes and genome annotation using RNA-seq identified the locations of 19,124 protein-coding genes. The four short-read individuals were aligned to our reference genome to investigate the demographic history of the two species. A Principal Component Analysis clearly separated re-sequenced individuals, while inferring population size history using the Pairwise Sequentially Markovian Coalescent model noted an approximate species split one million years ago, and a single, possibly recently introgressed individual.

## INTRODUCTION

Hybridization and introgression can occur between closely related species brought into secondary contact (Chown *et al*. 2015). Hybridization can be an evolutionary dead end, or it can lead to adaptive introgression (Arnold and Martin, 2009; Abbott *et al*. 2013). Introgression can result in the merging of hybridizing forms, reinforcement of reproductive barriers through selection for assortative mating, and a non-neutral shift in fitness among introgressed individuals. In some instances, this enables the expansion of the introgressed species into a novel habitat (Arnold, 1992). Further complicating this, adaptive introgression combined with climate change can weaken reproductive isolation (Owens & Samuk, 2020). In its extreme form, hybridization can drive extinction through introgression (Rhymer & Simberloff, 1996).

An increase in global surface temperatures has led to range shifts among a variety of taxa on a global scale (Chen *et al*. 2011) and increasing secondary contact between closely related species (Krosby *et al*. 2015), leading to increased opportunities for hybridization (Garroway *et al*. 2010; Chunco, 2014). Climate-driven range expansions have been noted in mammals, insects, and fish (Moritz *et al*. 2008; Garroway *et al*. 2010; Muhlfeld *et al*. 2014; Scriber, 2014), among other taxa. Instances of hybridization in wild ecosystems can be exacerbated by climate change because of increased secondary contact, where barriers to interspecific reproduction are reduced or removed altogether (Chunco, 2014).Without such barriers, species that were previously allopatric might interbreed, possibly leading to genetic admixture and potentially outbreeding depression or heterosis (Barton, 2001; Rius & Darling, 2014).

As climate-mediated range expansion has been shown to increase distributional overlap between related species (Chunco, 2014), climate change will therefore likely drive interspecific hybridization between many taxa. For example, studies in North America have noted hybrid zones across a latitudinal gradient between southern (*Glaucomys volans*) and northern (*Glaucomys sabrinus*) flying squirrels (Garroway *et al*. 2010; Rogic *et al*. 2016). Interspecific hybridization is a key conservation concern for these flying squirrel species, as population declines among northern flying squirrels have been noted in some areas of the USA, where some populations are endangered (Wood *et al*. 2016). The potential for introgressive hybridization and the subsequent ecological and fitness consequences necessitates a holistic assessment of species biology in the *Glaucomys* hybrid zone. The hybrid zone is a valuable study system to facilitate the assessment of interspecific hybridization, the potential for reinforcement of reproductive barriers, and the associated ecological conclusions in a wild, in-vivo system.

Low hybrid fitness can also lead to increased divergence between species through reinforcement. *Glaucomys* hybrid offspring are viable and can successfully backcross with either parental species (Garroway *et al*. 2010), however, the fitness implications among hybrid or introgressed individuals is unknown. The purpose of our study was to generate a *de novo* reference genome for *Glaucomys*. We annotated the reference genome using our already assembled and annotated flying squirrel transcriptome (Brown *et al*. 2021). Subsequently, using short reads from four individuals, two northern and two southern flying squirrels, we assembled re-sequenced high coverage genomes by aligning to the reference genome for a comparative analysis and demographic history reconstruction.

## MATERIALS AND METHODS

### Sample preparation

We isolated brain tissue from two adult *Glaucomys volans* and two adult *Glaucomys sabrinus* for sequencing. *G. sabrinus* individuals were collected from near Kawartha Highlands Signature Site Park (NFS_6525) and in Algonquin Provincial Park, Ontario, Canada (NFS_50254), and *G. volans* individuals were sampled near Sherborne Lake (SFS_25428) and Clear Creek, Ontario, Canada (SFS_CC1; Fig. 1). Algonquin Provincial Park (NFS_50254) was outside the northern edge of the hybrid zone, and Clear Creek (SFS_CC1) was outside the range of *G. sabrinus* and not an area of sympatry. The sites were all mature, closed canopy forest with a mixture of temperature deciduous trees such as sugar maple (*Acer saccharum*), red oak (*Quercus rubra*), and American beech (*Fagus grandifolia*), and coniferous trees such as white pine (*Pinus strobus*) in uplands or white spruce (*Picea glauca*) and balsam fir (*Abies balsamea* in riparian areas (see Bowman *et al*. 2005 for more details). All four specimens were morphologically identified to their parental species. Squirrel tissue samples were extracted using an organic extraction. The extracted DNA was run on a 1.5% agarose gel and Qubit fluorometer using the High Sensitivity Assay Kit to ensure we had sufficient DNA. They were also run on a Nanodrop ND-8000 spectrophotometer to test purity. The DNA was normalized to 20ng/μl at a final volume of 50μl.

**Figure 1.**
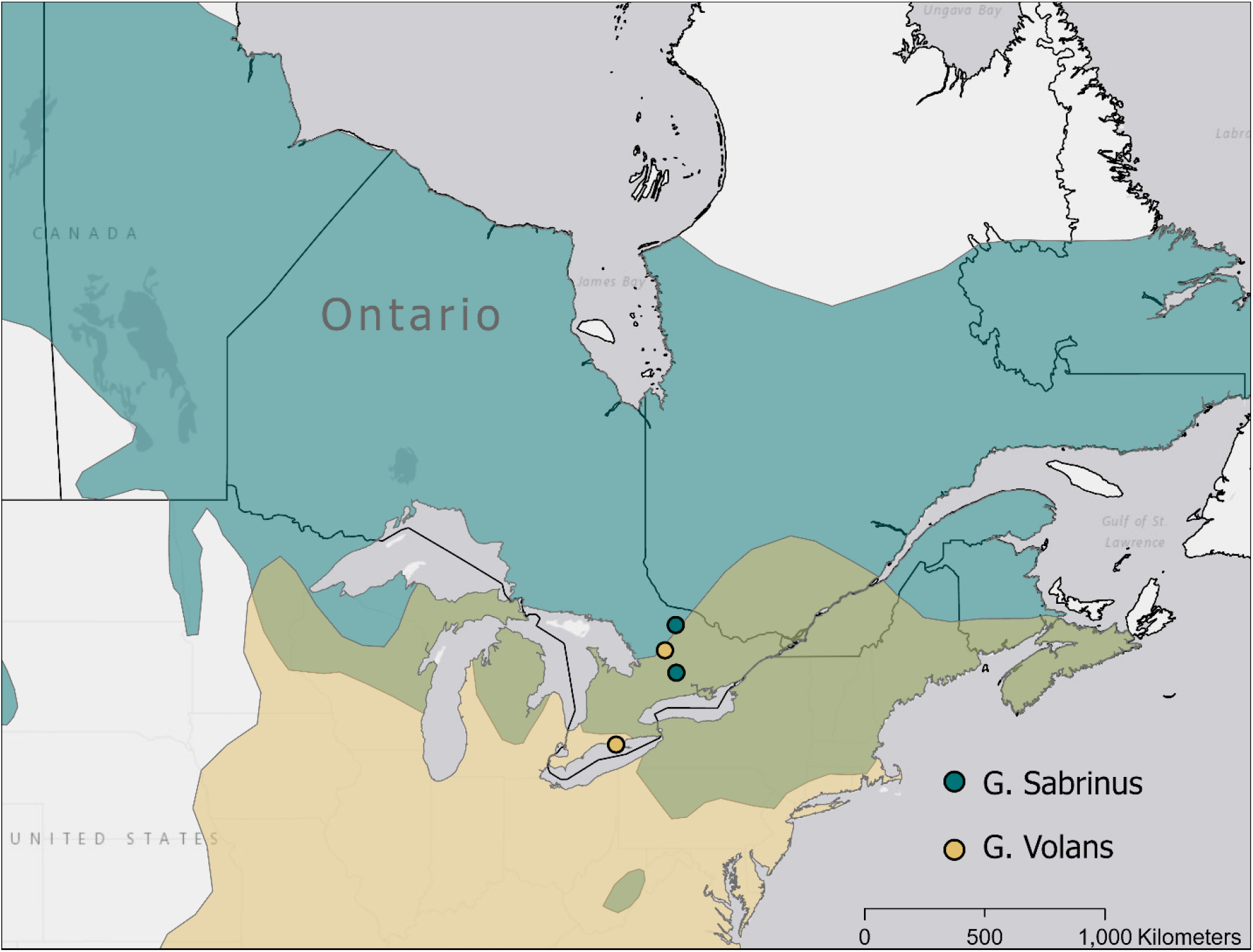
Range of northern (*Glaucomys sabrinus*) and southern (*Glaucomys volans*) flying squirrels overlaid with sampling locations. The geographic ranges are represented in the same colors as samples, while the hybrid zone is represented in olive, and only the southernmost *G. volans* sample from Clear Creek, Ontario, is located outside of the hybrid zone.

### *de novo* genome assembly

Southern flying squirrel libraries from individual CC1 were prepared and paired-end sequenced on 1 lane on an Illumina HiSeq X to generate 150 base pair (bp) paired-end reads. Sequencing was conducted at The Centre for Applied Genomics (Next Generation Sequencing Facility, SickKids Hospital, Toronto, Ontario, Canada). The sequence reads from each sample were provided in a FASTQ file format. 10X Genomics long read Chromium sequencing was used to generate linked reads. The estimated genome size was thought to be near that of the giant flying squirrel (*Petaurista leucogenys*) genome (C-value = 4.02 pg; Gregory, 2005). We used FastQC (version 0.11.9; Andrews, 2010) to perform simple quality control checks on raw sequence data to confirm the quality of the trimmed sequence reads. Long reads were assembled using Supernova as this assembler uses 10X linked-reads to produce phased assemblies of homologous chromosomes over multi-megabase ranges (Weisenfeld *et al*. 2018). Supernova recommends against trimming; however, Supernova was run using all available reads and was performed on both untrimmed and trimmed reads, to ascertain the impact of trimming on summary statistics. Trimming was completed using Trimmomatic v0.39 with two different parameter specifications as follows: 1) Illumina adapters were removed, leading and trailing low quality or N bases were removed (below quality 3), reads were scanned with a 4-base sliding window and cut when the average per quality base drops below 15, and reads were dropped that were less than 36 bases long after the previous steps and 2) Illumina adapters were removed (Bolger *et al*. 2014). Running Supernova on trimmed reads and parameter set 1) resulted in decreased raw coverage and Supernova was unable to generate an assembly. Running Supernova on trimmed reads and parameter set 2) resulted in an assembly with a slightly lower contig and scaffold N50 and thus, all subsequent analyses used the untrimmed read assembly. The FASTA file representing the assembly was generated using the pseudohap style output as it was the most contiguous. Assembly statistics were generated using BBMap 38.90 (Bushnell *et al*. 2017). However, scaffold N50 values were extremely similar regardless of the style of output that was selected. We used BUSCO (Benchmarking Universal Single-Copy Orthologs; (Waterhouse *et al*. 2018)) to reconstruct 4,104 conserved mammalian genes to assess genome completeness.

### Re-sequenced genome assemblies

Northern and southern flying squirrel libraries were prepared and paired-end sequenced across 8 lanes on an Illumina HiSeq X to generate 150 base pair (bp) paired-end reads. Sequencing was conducted at The Centre for Applied Genomics (Next Generation Sequencing Facility, SickKids Hospital, Toronto, Ontario, Canada). Forward and reverse reads were concatenated across eight lanes. FastQC was run as above to determine forward and reverse read quality and inform subsequent trimming parameters. We trimmed the adapters and low-quality bases from the reads with Trimmomatic as per the parameter set 1) mentioned above. To avoid any potential contamination of the genome sequence with viral or bacterial sequences, we screened the trimmed reads with Kraken2 (Wood *et al*. 2019) using the full standard database.

Reads from four individuals (NFS_6525, NFS_50254, SFS_25428, SFS_CC1) were aligned to the long read reference genome using Bowtie2 2.2.4 (Langmead & Salzberg, 2012), and the SAM file converted to a BAM file using Samtools 1.7 (Li *et al*. 2009). We removed poorly mapped reads via skipping alignments with MAPQ values smaller than 20 using Samtools 1.7. We removed duplicate reads and added correct read group information to each BAM file using Picard 2.18.27 (http://broadinstitute.github.io/picard/). We then clipped overlapping regions using clipOverlap from bamUtil 1.0.1.4 (Jun *et al*. 2015) and sorted the BAM file using Samtools 1.7 and built an index using Picard. All BAM files were checked using FastQC 0.11.9 (Andrews, 2010), and we calculated the mean depth of coverage for each BAM file using Samtools. We used Haplotype Caller in gatk 3.8 (Mckenna *et al*. 2010) to call variants and produce a variant call format (VCF) file for each flying squirrel. Individual VCF files were combined using the Combine GVCFs function, and then, we performed joint genotyping using Genotype GVCFs, both in GATK, to produce a VCF file with both northern and southern flying squirrels. We did some additional filtering on the combined VCF files to ensure quality. We used VCFtools 0.1.16 (Danecek *et al*. 2011) to do two rounds of filtering. First, we removed indels (using the remove-indels command), and any site with a depth of less than five or more than 33 (approximately double the average depth across the genome, using the min-meanDP and max-meanDP commands) and removed any low-quality genotype calls, with a score below 20, (using the minGQ command) which in VCFtools are changed to missing data. In the second round, we filtered to remove genotypes with more than 10% missing data (using the max-missing command). We did not filter to remove any SNP with a minor allele frequency (MAF) as we have only one or two individuals from each location and this results in removing the private sites, instead relying on very high depth and stringent filtering to ensure a high-quality data set.

The combined VCF file used for analyses with all individuals contained 35,937,561 SNPs. After filtering, we measured the mean depth (using the depth command) and the frequency of missing data (using the missing-indv command) for each individual in the final VCF file of 2 northern and 2 southern flying squirrels using VCFtools.

### Annotation

We identified and classified the repeat regions of the assembled genome using RepeatMasker v. 4.1.0 (Smit *et al*. 2013). We configured RepeatMasker with RMBlast v. 2.10.0 sequence search engine, Tandem Repeat Finder v. 4.0.9 (Benson, 1999), Dfam_Consensus database 3.1 (November 2020 release), and used the ‘-species glaucomys’ parameter for the analysis.

We used the gene prediction program AUGUSTUS 2.5.5 (Hoff & Stanke, 2019) to annotate the masked genome using predictions based on human genes. Additionally, we incorporated RNA-seq data into AUGUSTUS using the transcriptome created by Brown *et al*. (2021). We used BLAT v. 1.04 to help identify exon structure and allow for the subsequent generation of both intron and exon hints from alignments for AUGUSTUS (Hoff & Stanke, 2019; http://augustus.gobics.de/binaries/readme.rnaseq.html). The genome run in AUGUSTUS used a partial gene model allowing the prediction of incomplete genes at the sequence boundaries. The masked genome was split into 31 parts of ~1995 sequences each to reduce the computational resources and we concatenated the 31 output General Feature Format (GFF) files into a single annotation file.

### Comparative analyses

To compare whole-genome heterozygosity estimates, we used ANGSD to generate a site frequency spectrum and obtain heterozygosity values for each individual. We used the parameters -C 50 -ref ref.fa -minQ 20 -minmapq 30 to remove the low-quality bases and reads (Korneliussen *et al*. 2014). We generated a Principal Component Analysis (PCA) to determine if northern and southern flying squirrels grouped together or separately. We also ran Pairwise Sequentially Markovian Coalescent (PSMC; https://github.com/lh3/psmc) to model the historical effective population size and reconstruct the demographic history of both our northern and southern flying squirrel genomes. We used the default parameters of 64 atomic time intervals (-p “4+25*2+4+6”), a generation time of 1.5 years (COSEWIC, 1998), and a mutation rate of m = 2.0*10^-9^ mutations/site/generation (Gossmann *et al*. 2019).

## RESULTS AND DISCUSSION

### *G. volans* genome assembly

The final *Glaucomys volans* genome assembly was the untrimmed linked-read 10X Chromium assembly with Supernova (Weisenfeld *et al*. 2018), which produced a genome consisting of 7,087 scaffolds ≥50Kb with a scaffold N50 of 455.26Kb, a contig N50 of 75.63Kb, a GC content of 40.48%, and a genome size of 2.40Gb (Table 1; Table 2). These values were slightly more contiguous relative to a trimmed linked-read 10X Chromium assembly with Supernova (N50 = 450.38Kb, contig N50 = 78.87Kb, genome size = 2.39Gb). BUSCO indicated the presence of 3,742 (91.2%) complete mammalian genes of the 4,104 searched for. Our estimated genome size was similar to the assembly of the thirteen-lined ground squirrel (*Ictidomys tridecemlineatus*; ~2.5Gb), whereas the BUSCO value for the ground squirrel was 92.9% (Di Palma *et al*. 2011). Genome annotation of our final genome incorporating RNA-Seq data identified the locations of 19,124 protein-coding genes compared to 28,262 protein-coding genes without using RNA-Seq data.

**Table 1.** Summary statistics of the long read *Glaucomys volans* reference genome

| Statistic | *Glaucomys volans* genome |
| --- | --- |
| Scaffold sequence total (bp) | 2.58 x 10<sup>9</sup> |
| Number of scaffolds | 61,815 |
| Scaffold N50 (bp) | 455,262 |
| Scaffold L50 | 1,582 |
| Scaffold N90 (bp) | 117,214 |
| Scaffold L90 | 5,080 |
| Contig sequence total (bp) | 2.53 x 10<sup>9</sup> |
| Number of contigs | 115,069 |
| Contig N50 (bp) | 75,631 |
| Contig L50 | 9,446 |
| Contig N90 (bp) | 21,155 |
| Contig L90 | 30,374 |

**Table 2.** Nucleotide base composition of the long read *Glaucomys volans* reference genome

| A | C | T | G | N |
| --- | --- | --- | --- | --- |
| 29.77% | 20.24% | 29.75% | 20.24% | 0.17% |

### Re-sequenced genome assembly

Trimming the concatenated short read pairs resulted in the removal of an average of 4.37% of reads. The human library was removed from the full standard database, as its inclusion resulted in a relatively high percentage reads mapped to human due to orthologous mammal genes. After removing the human library, 0.25-0.35% of the reads were classified as belonging to an identified bacterial taxon; screening trimmed concatenated short read pairs for bacterial contaminants resulted in the further removal of an average of 0.29% of reads. The final short read coverage for each of the four individuals were as follows: SFS_CC1 = 15.75X, SFS_25428 = 17.55X, NFS_50254 = 17.88X, NFS_6525 = 14.96X. Our final VCF file contained 10% missing data. For all individuals, observed heterozygosity exceeded expected, while inbreeding coefficients ranged from 0.002610402 – 0.003583892 (NFS_50254 = 0.002763883, NFS_6525 = 0.002610402, SFS_CC1 = 0.003111787, SFS_25428 = 0.003583892).

### Comparative analyses and population history of *G. sabrinus and G. volans*

Northern and southern flying squirrels grouped distinctly in our PCA, while there was more variation among southern flying squirrels (Fig. 2). The first principal component accounted for over 80% of the variation noted, and clearly separated both species. Both southern individuals had higher whole-genome heterozygosity relative to northern individuals. There are multiple possible explanations for this result. For example, southern flying squirrels are smaller-bodied and typically exhibit higher population sizes and densities, whereas a lower effective population size in northern flying squirrels may result in decreased heterozygosity (Arbogast, 2007; Bowman *et al*. 2020). Overall, the levels of heterozygosity of both flying squirrel species are comparable to other genome-wide estimates in mammals (see Fig. 3 in Morin *et al*. 2021).

**Figure 2.**
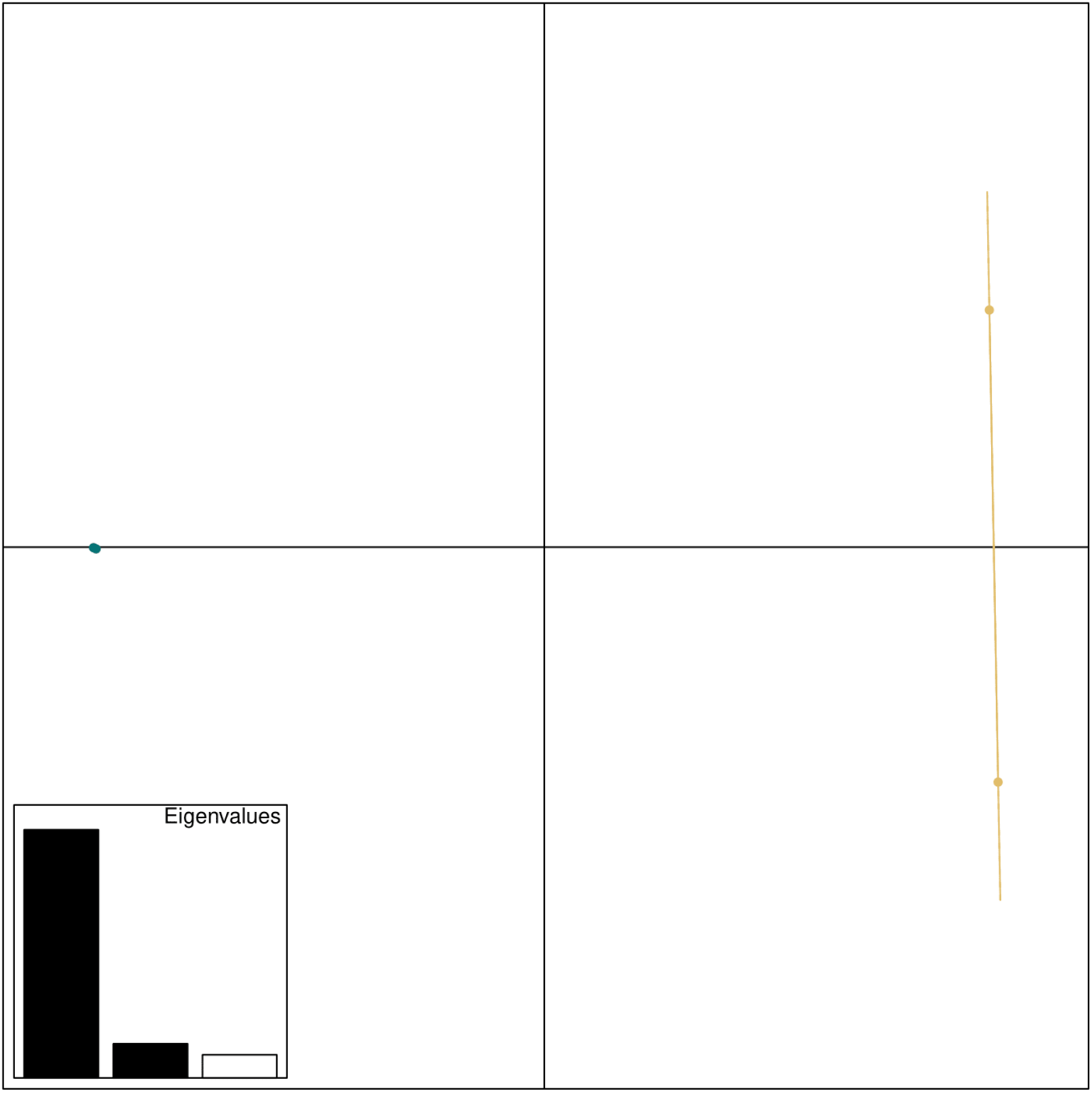
Principal Component Analysis (PCA) of 2 northern (*Glaucomys sabrinus* – represented in turquoise) and 2 southern (*Glaucomys volans* – represented in yellow) flying squirrel genomic variation. PC1 (x-axis) accounts for 81.23% of the variation, while PC2 (y-axis) accounts for 11.19% of the variation; the first two principal components account for over 90% of the genomic variation.

**Figure 3.**
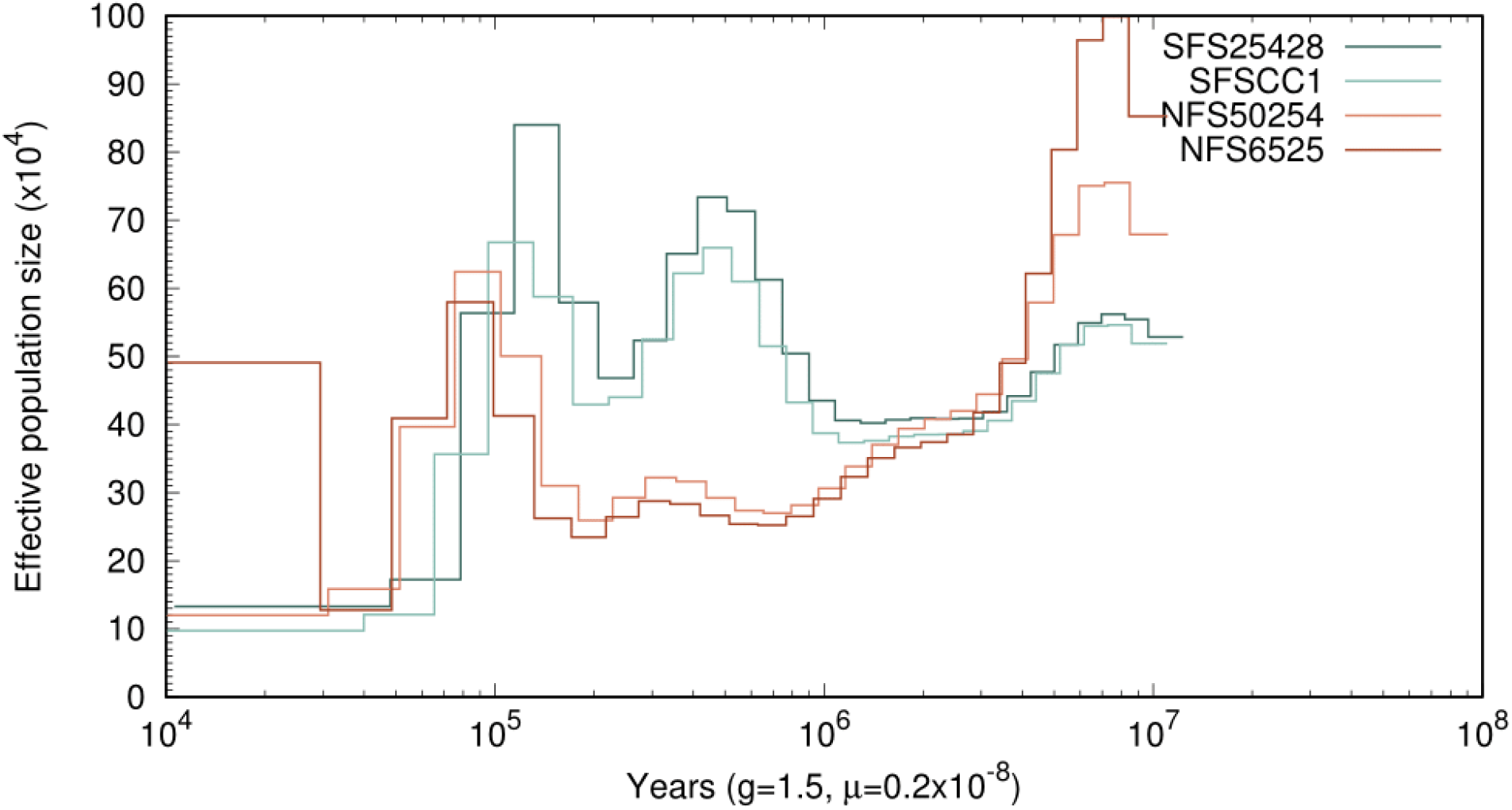
Reconstruction of historical effective population size (N_e_) of both northern (*Glacuomys sabrinus*) and southern (*Glaucomys volans*) flying squirrels using PSMC analysis assuming a mutation rate μ of 2.0 x 10^-9^ mutations/site/generation and a generation time of 1.5 years. Ne is in units of 10,000 individuals on the y-axis and time on the x-axis.

Previous research has estimated the split between northern and southern flying squirrels to be in the early to mid-Pleistocene (2,580,000 to 130,000 years ago; Arbogast, 1999, 2007). Based on PSMC analysis, the split between the species seemed to occur approximately 1mya, whereas, after 1mya, the species exhibited different trajectories (Fig. 3). Additionally, it is interesting that NFS_6525 had an increase in effective population size more recently, but NFS_50254 did not (Fig. 3). Previous work using microsatellites has been consistent with panmixia in Ontario within each of these species (Garroway *et al*. 2011; Bowman *et al*. 2020). It is possible however, that long-term introgression is evident in the genome of this northern flying squirrel individual (NFS_50254), leading it to group more closely to the southern flying squirrels. Analysis of a larger sample of genomes with varying degrees of introgression will help to clarify these patterns.

## CONCLUSION

High Throughput Sequencing studies on hybrid zones of wild non-model species have revealed traits associated with divergence in sympatry and allopatry (Scordato *et al*. 2017), patterns of introgression that differ between populations (Nolte *et al*. 2009), and genes associated with reproductive isolation (Teeter *et al*. 2008). Whole genome sequencing provides insight into the evolutionary process of hybridization and adaptive introgression, however, demonstrating the adaptive or fitness values of introgressed genomic regions remains an area of difficulty (Taylor & Larson, 2019). Studies of this kind benefit from a reference genome as a basis for identifying genomic regions of interest, and against which it is possible to evaluate potential hybrids and introgressed individuals (Payseur & Rieseberg, 2016).

As such, we produced a high-quality southern flying squirrel reference genome, an annotation in gff3 and bed format, and a RepeatMasked version of the genome, as well as high-coverage northern and southern flying squirrel re-sequenced genomes. The availability of a high-quality reference genome is invaluable in answering evolutionary questions surrounding hybridization and introgression and for conservation efforts. This is the first flying squirrel genome generated and will help future research determine not only the presence of hybrids in the North American flying squirrel hybrid zone but can also aid in identifying loci of interest in these same populations.

## DATA AVAILABILITY

10X Chromium long-read and Illumina short read data as well as a FASTA file of the assembly are available at the National Centre for Biotechnology Information (NCBI), under the BioProject accession number PRJNA723586. The Whole Genome Assembly has been deposited at NCBI under the BioProject accession number PRJNA723289; BioSample number SAMN18810840.

## ACKNOWLEDGEMENTS

This research was supported by NSERC Discovery Grants to JB and PJW, and by the Ontario Ministry of Natural Resources and Forestry. The authors would like to thank: Michael G.C. Brown for assistance with sampling. Bridget Redquest and Austin Thompson for DNA extractions, and Kathleen Lo for their comments on earlier drafts of the manuscript.

## LITERATURE CITED

Abbott, R., D. Albach, S. Ansell, J. W. Arntzen, S. J. E. Baird et al. 2013 Hybridization and speciation. J. Evol. Biol. 26: 229–246.

Andrews, S. 2010. FastQC: a quality control tool for high throughput sequence data. Available online at: http://www.bioinformatics.babraham.ac.uk/projects/fastqc

Arbogast, B. S., 2007 A brief history of the new world flying squirrels: Phylogeny, biogeography, and conservation genetics. J. Mammal. 88: 840–849.

Arbogast, B. S., 1999 Mitochondrial DNA phylogeography of the new world flying squirrels (Glaucomys): Implications for pleistocene biogeography. J. Mammal. 80: 142–155.

Arnold, M. L., 1992 Natural hybridization as an evolutionary process. Annu. Rev. Ecol. Syst. 23: 237–261.

Arnold, M. L., and N. H. Martin, 2009 Adaptation by introgression. J. Biol. 8: 9–11.

Barton, N. H., 2001 The role of hybridization in evolution. Mol. Ecol. 10: 551–568.

Bolger, A. M., M. Lohse, and B. Usadel, 2014 Trimmomatic: A flexible trimmer for Illumina sequence data. Bioinformatics 30: 2114–2120.

Bowman, J., P. O’Brien, and P. J. Wilson, 2020 Landscape genetics of flying squirrels in Ontario.

Bowman, J., G. L. Holloway, J. R. Malcolm, K. R. Middel, and P. J. Wilson. 2005 Northern range boundary dynamics of southern flying squirrels: evidence of an energetic bottleneck. Can. J. Zool. 83: 1486–1494.

Brown, M. G. C., J. Bowman, and P. J. Wilson, 2021 Novel de novo transcriptome assembly, functional annotation, and SNP discovery in North American flying squirrels (genus Glaucomys).

Bushnell, B., J. Rood, and E. Singer, 2017 BBMerge – Accurate paired shotgun read merging via overlap. PLoS One 12: 1–15.

Chen, I. C., J. K. Hill, R. Ohlemüller, D. B. Roy, and C. D. Thomas, 2011 Rapid range shifts of species associated with high levels of climate warming. Science (80-.). 333: 1024–1026.

Chown, S. L., K. A. Hodgins, P. C. Griffin, J. G. Oakeshott, M. Byrne et al. 2015 Biological invasions, climate change and genomics. Evol. Appl. 8: 23–46.

Chunco, A. J., 2014 Hybridization in a warmer world. Ecol. Evol. 4: 2019–2031.

COSEWIC. 1998. Southern flying squirrel (Glaucomys volans) COSEWIC assessment and status report. Available from: https://www.canada.ca/en/environment-climate-change/services/species-risk-public-registry/cosewic-assessments-status-reports/southern-flying-squirrel/chapter-2.html

Danecek, P., A. Auton, G. Abecasis, C. A. Albers, E. Banks et al. 2011 The variant call format and VCFtools. 27: 2156–2158.

Di Palma F, et al. 2011. The draft genome of *Spermophilus decemlineatus*. GenBank GCA_000236235.1

Garroway, C. J., J. Bowman, T. J. Cascaden, G. L. Holloway, C. G. Mahan et al. 2010 Climate change induced hybridization in flying squirrels. Glob. Chang. Biol. 16: 113–121.

Garroway, C. J., J. Bowman, G. L. Holloway, J. R. Malcolm, and P. J. Wilson. 2011 The genetic signature of rapid range expansion by flying squirrels in response to contemporary climate warming. Global Change Biol. 17: 1760–1769.

Garroway, C. J., J. Bowman, T. J. Cascaden, G. L. Holloway, C. G. Mahan, J. R. Malcolm, M. A. Steele, G. Turner, and P. J. Wilson. 2010 Climate change induced hybridization in flying squirrels. Global Change Biol. 16: 113–121.

Gossmann, T. I., A. Shanmugasundram, S. Börno, L. Duvaux, C. Lemaire et al. 2019 Ice-Age Climate Adaptations Trap the Alpine Marmot in a State of Low Genetic Diversity. Curr. Biol. 29: 1712–1720.e7.

Gregory, T.R. 2020. Animal Genome Size Database. http://www.genomesize.com.

Hoff, K. J., and M. Stanke, 2019 Predicting Genes in Single Genomes with AUGUSTUS. Curr. Protoc. Bioinforma. 65: 1–54.

Jun, G., M. K. Wing, G. R. Abecasis, and H. M. Kang, 2015 An efficient and scalable analysis framework for variant extraction and refinement from population scale DNA sequence data. Genome Res. gr-176552:

Korneliussen, T.S., Albrechtsen, A. & Nielsen, R. ANGSD: Analysis of Next Generation Sequencing Data. BMC Bioinformatics 15, 356 (2014). https://doi.org/10.1186/s12859-014-0356-4

Krosby, M., C. B. Wilsey, J. L. McGuire, J. M. Duggan, T. M. Nogeire et al. 2015 Climate-induced range overlap among closely related species. Nat. Clim. Chang. 5: 883–886.

Langmead, B., and S. L. Salzberg, 2012 Fast gapped-read alignment with Bowtie 2. 9: 357–360.

Li, H., B. Handsaker, A. Wysoker, T. Fennell, J. Ruan et al. 2009 The Sequence Alignment / Map format and SAMtools. 25: 2078–2079.

Mckenna, A., M. Hanna, E. Banks, A. Sivachenko, K. Cibulskis et al. 2010 The Genome Analysis Toolkit: A MapReduce framework for analyzing next-generation DNA sequencing data. 1297–1303.

Morin, P. A., F. I. Archer, C. D. Avila, J. R. Balacco, Y. V. Bukhman et al,. Reference genome and demographic history of the most endangered marine mammal, the vaquita. Mol. Ecol. Resour. 21: 1008–1020.

Moritz, C., J. L. Patton, C. J. Conroy, J. L. Parra, G. C. White et al. 2008 Impact of a century of climate change on small-mammal communities in Yosemite National Park, USA. Science (80-.). 322: 261–264.

Muhlfeld, C. C., R. P. Kovach, L. A. Jones, R. Al-Chokhachy, M. C. Boyer et al. 2014 Invasive hybridization in a threatened species is accelerated by climate change. Nat. Clim. Chang. 4: 620–624.

Nolte, A. W., Z. Gompert, and C. A. Buerkle, 2009 Variable patterns of introgression in two sculpin hybrid zones suggest that genomic isolation differs among populations. Mol. Ecol. 18: 2615–2627.

Owens, G. L., and K. Samuk, 2020 Adaptive introgression during environmental change can weaken reproductive isolation. Nat. Clim. Chang. 10: 58–62.

Payseur, B., and L. Rieseberg, 2016 A genomic perspective on hybridization and speciation Bret. Mol. Ecol. 25: 2337–2360.

Rhymer, J. M., and D. Simberloff, 1996 Extinction by hybridization and introgression. Annu. Rev. Ecol. Syst. 27: 83–109.

Rius, M., and J. A. Darling, 2014 How important is intraspecific genetic admixture to the success of colonising populations? Trends Ecol. Evol. 29: 233–242.

Rogic, A., G. Dubois, N. Tessier, P. Paré, P. Canac-Marquis et al. 2016 Applying genetic methods to identify northern and southern flying squirrels and determine conservation needs. Conserv. Genet. Resour. 8: 471–480.

Scordato, E. S. C., M. R. Wilkins, G. Semenov, A. S. Rubtsov, N. C. Kane et al. 2017 Genomic variation across two barn swallow hybrid zones reveals traits associated with divergence in sympatry and allopatry. Mol. Ecol. 26: 5676–5691.

Scriber, J. M., 2014 Climate-driven reshuffling of species and genes: Potential conservation roles for species translocations and recombinant hybrid genotypes.

Taylor, S. A., and E. L. Larson, 2019 Insights from genomes into the evolutionary importance and prevalence of hybridization in nature. Nat. Ecol. Evol. 3: 170–177.

Teeter, K. C., B. A. Payseur, L. W. Harris, M. A. Bakewell, L. M. Thibodeau et al. 2008 Genome-wide patterns of gene flow across a house mouse hybrid zone. Genome Res. 18: 67–76.

Waterhouse, R. M., M. Seppey, F. A. Simao, M. Manni, P. Ioannidis et al. 2018 BUSCO applications from quality assessments to gene prediction and phylogenomics. Mol. Biol. Evol. 35: 543–548.

Weisenfeld, N. I., V. Kumar, P. Shah, D. M. Church, and D. B. Jaffe, 2018 Direct determination of diploid genome sequences. Genome Res. 28: 757–767.

Wood, D. E., Lu, Jennifer, and B. Langmead, 2019 Improved metagenomic analysis with Kraken 2. bioRxiv 1–13.

Wood, C. M., J. W. Witham, and M. L. Hunter, 2016 Climate-driven range shifts are stochastic processes at a local level: Two flying squirrel species in Maine. Ecosphere 7: 1–9.

